# A dynamic inflammation model for neutrophil and monocyte responses in sepsis, trauma and surgery patient clusters

**DOI:** 10.1101/2023.02.14.527104

**Authors:** B. Browning, F. Couenne, C. Bordes, L. Fayolle, F. Venet, J. Textoris, G. Monneret, M. Tayakout-Fayolle

## Abstract

Time series clustering is applied to inflammation and neutrophil cell development markers, CD16 and CD10, in sepsis, trauma and surgery patients and a dynamical model with an inflammation function, *F*, is used to represent their evolution over a two month period. Five patient clusters are identified, characterised and evaluated against medical assessment scores and the literature. A dynamical model for neutrophil and monocyte cell counts and maturity has been constructed based on mass balances and cell kinetics in both blood and bone marrow. Cell proliferation and flow rates, as well as expression of monocyte HLA-DR, depend on concentrations of pro- and anti- inflammatory cytokines, IL6 and IL10, via *F*. A good fit with the data is obtained for each cluster and the estimated parameters correlate to illness severity. The model is a potential tool for simulation of immunomodulatory therapies.

## Introduction

Sepsis is a life-threatening organ dysfunction caused by a dysregulated host response to infection^1^. It has a huge global impact, with the number of deaths from sepsis similar to that for all cancers put together^2,3^. As such, it is a global health priority for the World Health Organisation^4^. Regarding pro-/anti-inflammatory overtime trajectories, septic patient groups are heterogeneous^5^. Thus, despite decades of active research^6–8^, anti-inflammatory medication, various immunomodulatory strategies and cytokine blocking therapies did not improve patient outcomes^9,10^. Importantly, recent results from the REALISM study demonstrated that there was no specific immune dysfunction to differentiate sepsis from trauma or surgery patients, indicating that a common injury-induced immune response exists each time the body faces tremendous systemic inflammation^5,11,12^.

Hotchkiss et al. hypothesised that, in sepsis, the proper balance between pro- and antiinflammatory pathways determines the fate of the individual. They describe a theoretical model beginning with infection. At the onset, cells of the innate immune system release high levels of pro-inflammatory cytokines that drive inflammation. This leads to a hyperinflammatory stage that can result in cytokine storm and death, but which most patients survive. If sepsis persists, the patient can enter an immunosuppressed state^13,14^ and this is the predominant driving force for morbidity and mortality. Sepsis advances rapidly and the immune system is strongly involved, with 60% of the patients genome altered within 30 minutes of hospital admission^15^ and cytokines and receptors linked to increased inflammation and innate immune response particularly affected^15^. IL6 and IL10 are pro- and antiinflammatory cytokines, that have been used as inflammation markers in several models^16–19^ and shown to increase production rates of altered neutrophils and monocytes in-vitro^20^.

Early IL6 concentrations have been found to predict post-traumatic complications^21^ whilst persistent high IL6 concentrations 3 days after injury predicted 28 day mortality in critically ill patients^22^. In dynamic modelling, Azhar et al.^16^ selected IL6 as biomarker for systemic inflammation level to model the control architecture for the response to traumatic injury and Namas et al.^18^ used IL6 and IL10 concentrations to calibrate their model of T-helper cell dynamics in acute inflammation. IL6 and IL10 circulate in healthy adults at levels from undetectably low to around 10 pg/mL ^11^. The principle function of IL10 is to limit, and ultimately terminate, inflammatory responses^23^. In this respect, IL10 is a wide acting and a potent moderator in the inflammatory response network. In particular, it inhibits its own production as well as that of interleukin, IL6^23^. IL6 and IL10 signal to cells by attaching to their surface via specific receptors. Tallon et al.^17^ used IL10 to represent the anti-inflammatory signal in their model of pro- and anti-inflammatory cytokines in the early stages of septic shock.

Neutrophils have evolved to coordinate inflammation in the body and send out signals by secreting chemokines^24^ and cytokines^24^, such as IL6 and IL10. They are the most abundant form of white blood cell^24^. Key components of the innate immune response, in healthy individuals they have a lifespan of around 24 hours^25^ and their release from the bone marrow is tightly regulated. They circulate in the blood stream but are also found in discrete intravascular marginated pools^26^. They seek out infection and inflammation and are lethal to invaders. Neutrophil count and maturity are intimately linked with sepsis severity. Neutrophil production and maturation occur within the bone marrow niches ^27,28^ with no macro-directional trend^29^. Neutrophils spend about six days from inception to release into the blood stream^26^. Low levels of cell development markers, CD16 and CD10, have been linked to deterioration in sepsis^30^. These are acquired during the maturation process^27,31^ with CD16 addition earlier than that of CD10^27,31^. Adimy et al.^32^ review model types for cell formation dynamics in the bone marrow. Recently, Mika et al^33^ used a compartment type model to represent neutrophil production whilst Cassidy et al.^34^ use a system of discrete delay differential equations to model chemotherapy induced neutropenia and monocytopenia. Orr et al.^31^ used a pipeline model to study the acquisition of CD10 expression.

Like neutrophils, monocytes are produced and mature in the bone marrow^34,35^. Once in circulation, monocytes continue to evolve slowly from classical (85%) to intermediate and non-classical form^36^. They also have plasticity and adapt under septic conditions^14^. Delay differential equations have been used to represent the kinetics of formation, maturation and lifetime of classical monocytes ^36,37^ with approximately 2 days estimated for both the maturation period and the lifetime in the circulation^36,37^. Human leukocyte antigen – DR (HLA-DR) is a protein expressed on monocytes in the bone marrow towards the end of the maturation period^38^. HLA-DR is vital for correct monocyte function^39^. Its role is to present peptides derived from ingested microbes to T-cells and thus initiate a specific immune response^39^ and its expression is usually increased during bacterial infection^40^. However, it is rapidly down-regulated in sepsis^39,41,42^ and surgery^43^ and low monocyte HLA-DR (mHLA-DR) is currently the best indicator for sepsis patients who might benefit from immunostimulatory treatment^44^. A key difficulty is that immunostimulatory treatment could be harmful to some sepsis patients. Tailored therapies are being developed to overcome this problem but this approach is still in its infancy^44^.

Clustering methods are one way to try and solve this issue. They have been applied to the evolution of sepsis in very large studies^45^ but also on the scale of a few hundred patients^5^. Leijte et al.^46^ carried out a detailed analysis of mHLA-DR expression kinetics in a large cohort of 241 septic shock patients over the first week after injury. Unsupervised cluster analysis showed that mHLA-DR expression kinetics were not related to the site of initial infection. They identified three mHLA-DR trajectories that corresponded strongly to patients outcomes: early improvers, delayed improvers and decliners. Bodinier et al.^47^ performed time series clustering on mHLA-DR data for two large cohorts of sepsis patients (n = 333 and 107) using the K-means for longitudinal data R-package. They confirmed the three endotypes found by Leijte et al.^46^ and identified a fourth group of ‘high expressors’, who recovered relatively quickly.

Dynamic modelling can also help differentiate patient groups because the early evolution of inflammation biomarkers gives a good measure of immune system health^13^. Foteinou et al.^48^ discussed a systems based approach to inflammation modelling based on transcription factor networks and reference IL6 and IL10 as important parts of the pro- and anti-inflammatory processes. Reynolds et al.^49^ constructed a dynamic model of the acute inflammatory response which has been used or adapted by several authors^19,50,51^. Day et al.^19^ used four differential equations to create a dynamic model for COVID19 where the variables were: number of virus infected cells, damage/dysfunction, adaptive immune response and innate immune response (inflammation). They identified four distinct dynamic patterns of viral load and immune response. In these models, inflammation severity is an abstract concept which cannot be measured^50^. Tallon et al.^17^ tested a dynamical model for pro- and anti- inflammatory cytokines in the early stages of septic shock. Their model was composed of mass balances on cytokines, receptors and white blood cells and included production, transfer, adsorption and decay. Cytokine production rates depended on adsorbed cytokine concentrations and cell production rates were calculated using an inflammation function. They estimated model parameters against measured white blood cell counts and transcriptomic data in 28 patients where blood samples were taken 6 hourly during the first 48h of septic shock. This gave a good fit with the model most sensitive to parameters associated with pro-inflammatory cytokines, in line with the idea that early sepsis is driven by inflammatory factors.

The objective here is to identify patient trajectories under different severities of trauma, surgery and sepsis. The aim is earlier identification of patients who would benefit from targeted treatments and, also, better understanding of the differences between a healthy and pathological response. We use time series clustering^52^ and a dynamical model, based on chemical engineering principals^53^, with data from the REALISM study^12^. We identify five patient groups with different recovery profiles after sepsis, trauma or surgery then represent their IL6 and IL10 trajectories as proxies for inflammatory and anti-inflammatory effects via an inflammation function^17^, *F*. This serves as input to a dynamical model which calculates neutrophil and monocyte counts and maturity characteristics.

### Database and methods

#### REALISM Database

Details of the REALISM study are reported elsewhere^12^. It includes comprehensive data from 353 sepsis, trauma and surgery patients, who were followed for two months after injury, plus matching data for 175 healthy volunteers. For each patient, up to seven blood samples were taken after injury. These were on the first day, D1, then, on days 2, 3-4, 5-7, 13-18, 26-36 and 52-68. Surgery patients were also tested beforehand on day zero, D0. Neutrophil and monocyte cell counts are available, as is quantitative data for mHLA-DR, reported in number of antibodies per cell (Ab/c), as well as blood concentrations for soluble IL6 and IL10. Neutrophils are classified into four groups by level of phenotype markers CD10 and CD16 as high (+) or low (-). Data from the REALISM study has already been used to highlight the link between profound immune alterations and secondary infections^11^ and in work to identify different patient endotypes by clustering mHLA-DR trajectories^46,47^.

### Database exploitation

#### Patient clustering

The first step of clustering was to review the data. The variation in total neutrophil count, the four neutrophil phenotypes plus IL6 and IL10 concentrations could be clearly observed across all the patients and these seven variables were selected for clustering. The data has three dimensions: time, patient identity and measurement. There is a very rapid initial dynamic and the start time is not precisely known for sepsis and trauma patients. This can mask the true pattern in the data because measured curves overlap. The tslearn^52^ time series K-means package with the dynamic time warping metric is designed to overcome this and was used for clustering in Python with sci-kit learn^54^. MinMaxScaler^54^ was used to adjust the data to the same scale and natural logs were taken of the IL6 and IL10 concentrations to avoid ‘crushing’ the data. The clustering package requires that each measure has a value for every timepoint. To maximise use of the data with minimum interpolation, measurements from D1 to D18 were used as clustering input. In total, this comprised 8768 data points from 1270 blood samples and 353 patients. For each patient, results for days with no sample were calculated by averaging timepoints directly before and after gaps. The inertia is the sum of the squares of the distances between each data point and its cluster centre and is generally used to empirically determine an appropriate number of clusters. From this, we selected five clusters as optimum.

#### Dynamical model description

Having defined the clusters, we represent them with a dynamical model based on chemical engineering principals^53^. The model is shown in Figure 1. It represents the evolution of neutrophils, monocytes and HLA-DR proteins through the bone marrow and into the circulation under the influence of an inflammation function, *F* ^17^. The model is comprised of mass balances and represents the bone marrow and blood circulation as a series of stirred tanks or compartments. Cell transfer occurs as liquid plasma flows through the theoretical tanks carrying neutrophils and monocytes. Given that no macro-directional trend is observed for maturation of neutrophils or monocytes in the bone marrow^29^, we assume uniform blood dispersion and distribution of the very small scale niches throughout the bone marrow and then sum these to get to the model. *F* derives from the cell behaviour but here it is calculated directly from the measured IL6 and IL10 concentrations. Proliferation occurs only in compartment 1. Cells mature in compartments 2 and 3 and cell death occurs only in the circulation. *F* acts on cell proliferation, plasma flow and mature neutrophil and mHLA-DR disappearance rates. The initial disappearance rates of CD16+ CD10+ neutrophils and mHLA-DR from the circulation are too rapid to be a consequence only of normal cell death (apoptosis) and so we chose to apply the inflammation function to them directly. An increase in vascular permeability is associated with inflammation^50^ and is represented by a leakage stream directly from each compartment to the circulation.

**Figure 1.**
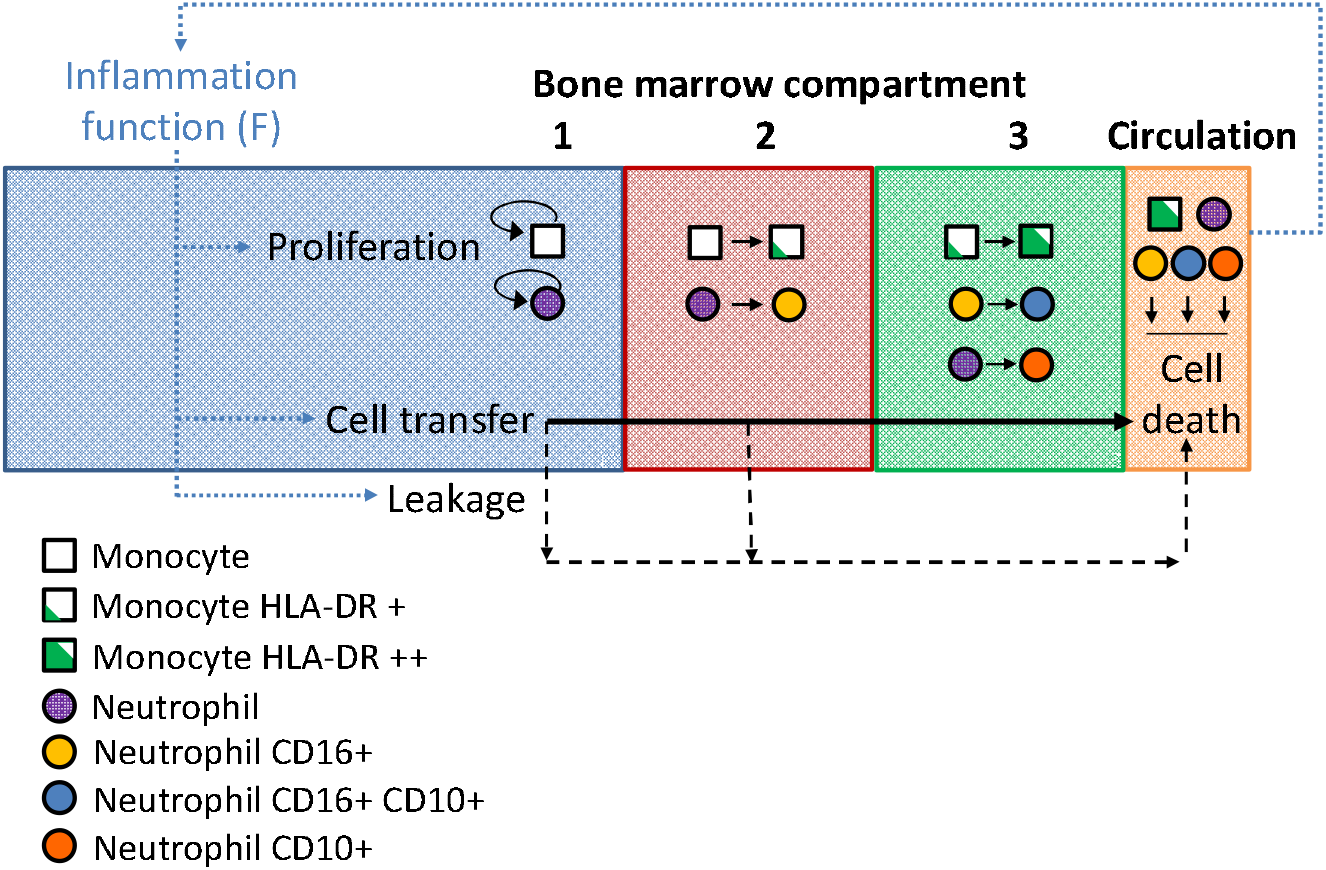
Flow diagram of neutrophil and monocyte model

The REALISM study measured neutrophil and monocyte maturity in different ways. Neutrophil maturity is indicated by thresholds for CD16 and CD10 acquisition, resulting in four phenotypes. Figure 2 shows the reaction scheme used in the model. In compartment 2, immature, CD10-CD16-, neutrophils are converted into CD10-CD16+ with the rate constant *k_nhl_*. In compartment 3, CD10-CD16+ and unconverted CD10-CD16-neutrophils evolve into CD10+ CD16+ and CD10+ CD16-respectively with reaction rate constants: *k_nhh_* and *k_nlh_*. Monocyte maturation is different in that the maturity measure, HLA-DR per monocyte, is continually increasing during the monocyte’s residence in compartments 2 and 3.

**Figure 2.**
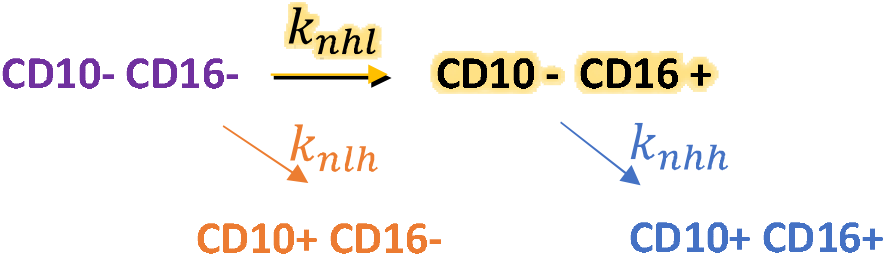
Reaction scheme for neutrophil maturation. CD16 is added in compartment 2 and CD10 is added in compartment 3.

The model requires estimates of each compartment volume. We assume the same compartments for monocytes and neutrophils. The distribution of neutrophils at different maturation stages and in circulation is not precisely known^26^ and varies between individuals^27^. Cell development marker, CD16 is added before CD10^31^ and about half the neutrophils in the body are undergoing proliferation or maturation pre-CD16 acquisition^26,27^. Most of the neutrophils are in the bone marrow with 5 – 15 % in the blood^26,33^. We assume cell distribution and compartment volumes shown in Table 1 based on a total bone marrow volume of 1.75L^26,27,29^.

**Table 1.** Distribution of cell quantities between compartments and compartment volumes.

| Compartment | 1 | 2 | 3 | Circulation |
| --- | --- | --- | --- | --- |
| Cell behaviour | Proliferation | Add CD16 / HLA-DR | Add CD10/ HLA-DR | Death |
| Cell fraction | 0.5 | 0.2 | 0.2 | 0.1 |
| Compartment volume (L) | 0.972 | 0.389 | 0.389 | 5 |

*F* is determined from Eq.s (1) and (2). Here, *C*_*IL*6_ and *C*_*IL*10_ are the molar blood concentrations of IL6 and IL10, proxies for pro- and anti- inflammatory effects in the body. Soluble IL6 and IL10 are converted from measured mass concentrations using their molar masses, respectively 21 and 18 kg/mol. The term, *α* = 0.848, is determined from the average measured values for healthy individuals, *C*_*IL*6,*h*_ and *C*_*IL*10,*h*_. In both bone marrow and circulation, the cell response to *F* is bound by physical limits. For example, surface receptor quantity and quality, i.e. affinity for associated cytokines^17^. To account for this we apply Eq. (3), based on the Langmuir function^55^. Parameters, *a* and *b*, are estimated where *a* represents receptor quantity and, *b*, the affinity. So, four inflammation responses, *f_n_, f_m_, f_H_* and *f_nhh_* are calculated in the model with two parameters estimated for neutrophils, monocytes and mHLA-DR: *a_n_, b_n_, a_m_, b_m_* and *a_H_, b_H_*. Finally, only one parameter was significant for the mature neutrophils, *b_nHH_* (see Nomenclature).

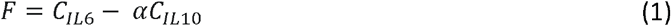

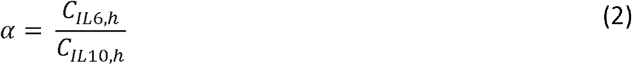

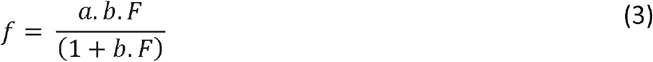

Neutrophil and monocyte proliferation occurs in bone marrow compartment 1. The relevant mass balances are given in Eq.s (4) and (5). *V* is the compartment volume in L. *C_N_* refers to the cell concentration in Giga/L, *nll* to immature neutrophils, *m* to monocytes and 1 to the compartment number. *k_p_* is the cell proliferation rate constant in 1/d, *Q* the liquid plasma flow rate in L/d, and *x* is a leakage factor. At steady state, the accumulation term and *f* are both zero and the cell outflow rate equals the production rate. The production and outflow rates increase with *f*, as does leakage directly from compartment 1 to the circulation.

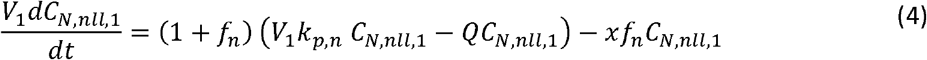

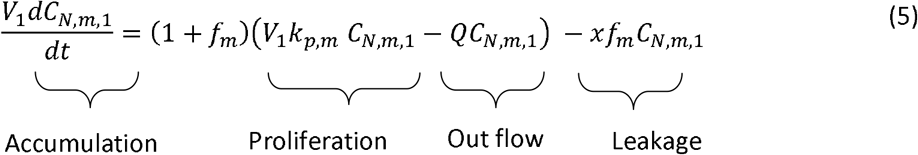

Neutrophils and monocytes mature in compartments 2 and 3. There are six mass balance equations for the neutrophils in compartments 2 and 3. The general equation is given in Eq. (6), where *i* is the compartment number. The neutrophil phenotype, *nll, nhl* or *nlh*, is represented by *nj*. The reaction, i.e. cell development marker and neutrophil phenotype combination, is denoted by *nr*. There are three possibilities as shown in Figure 2. *CD*16 can be added to *nll* and *CD*10 can be added to *nll* or *nhl*.

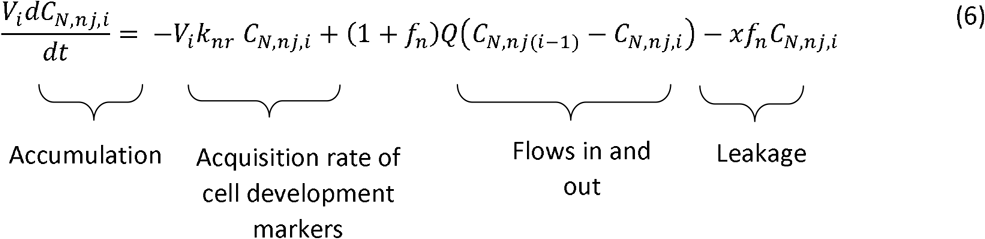

General mass balances for monocytes and HLA-DR in compartments 2 and 3 are given in Eq.s (7) and (8), where *N_H_* is the quantity of antibodies, Ab, in Ab/cell.

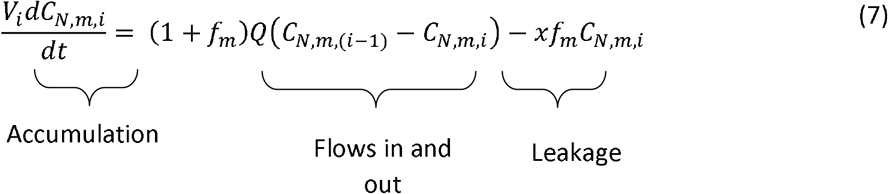

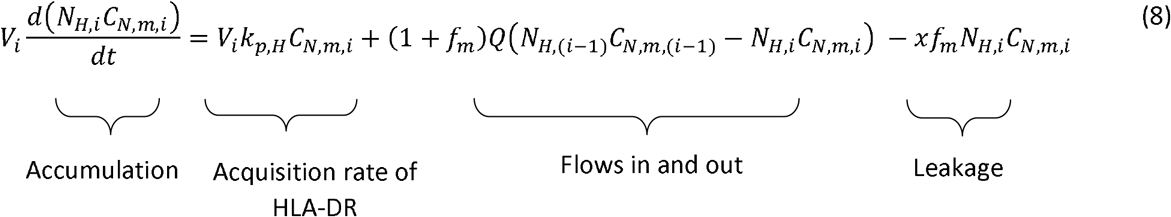

The general mass balances for cells and HLA-DR in circulation are given in Eq.s (9) and (10). In Eq. (9), *c* indicates circulation and *k_d_* is the disappearance rate constant in 1/d. The term, *f_nj_*, is set to zero except for mature, CD16+ CD10+, neutrophils.

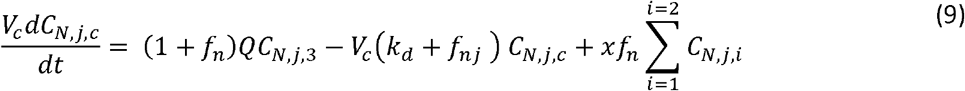

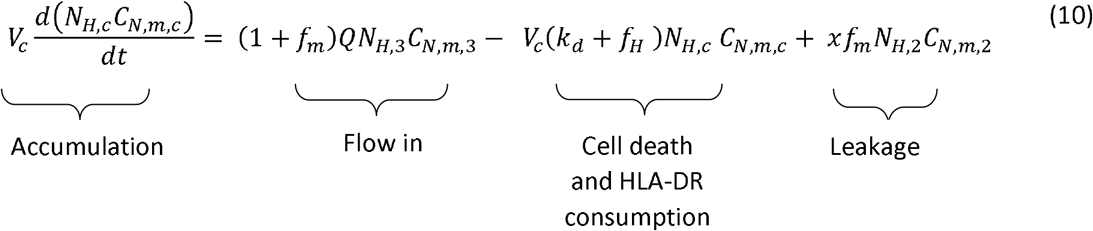

The system of differential equations was resolved in MATLAB R2019b^56^ using the solver ode15i and parameter estimation was carried out using MATLAB R2019b^56^ non-linear least squares solver function, lsqnonlin, was used with the trust-region-reflective algorithm. This minimises the objective function based on an input vector of differences between measured and calculated data. Parameter significance levels and confidence limits were determined from Eq.s (11) and (12)^57^. *J* is the Jacobian matrix and *se*(*b_p_*) is the standard error found for each parameter. *SSE, n_d_* and *p* are the objective function and the numbers of data points and parameters respectively. *b_p_* is the parameter value, *t_obs_* is the observed student t-value. The parameter significance levels are found from the Student t-table. Eq. (13) is the formula for the correlation between parameters *p* and *q*. For each cluster, parameter estimation was carried out against all the available data points from day 1 to 69 (with outliers removed). The data was weighted to balance the different measurement scales.

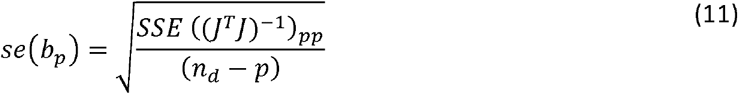

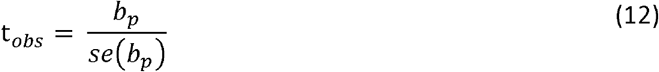

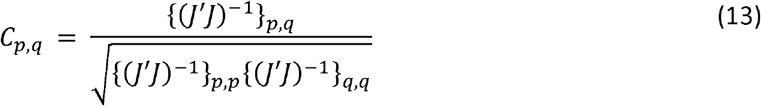

The measured IL6 and IL10 data is in discrete form, reported by day, whereas continuous IL6 and IL10 concentrations are needed for the model inputs. The ‘Dynamics of a Cytokine Storm’ model by Yiu et al.^58^ fit the first two days data well, but not a longer period, and the data was not smooth enough for interpolation. Consequently, we developed an in-house model for this.

Rate constants for neutrophil and monocyte death are set to 1 per day based on the literature^17^. So, a number of the parameters in Eq.s (4) to (10) are pre-determined. These are liquid plasma flow rate, *Q* = 0.194 L/d, and proliferation rate constants, *k_p,n_* = 0.2 d^−1^ and *k_p,m_* = 0.2 d^−1^. We assume that individuals have intrinsic constant neutrophil and monocyte maturation rates and each cluster includes surgery patients, so these constants can also be calculated. We treat the monocytes and neutrophils differently because we have more measurements for the neutrophils. For the monocytes we extrapolate the *k_p,H_* values to the whole cluster and use them in the model. For the neutrophils we set the calculated initial values of *k_nhl_, k_nhh_* and *k_nlh_*, based on the surgery patients, as the initial values for parameter estimation. Table 2 lists the parameters for estimation.

**Table 2.** Parameters for estimation

|  |  |  |
| --- | --- | --- |
| $a_n$ | For Langmuir function (neutrophils) | (-) |
| $b_n$ | For Langmuir function (neutrophils) | (-) |
| $k_{nhl}$ | Rate constant for cell development marker acquisition | (1/d) |
| $k_{nlh}$ | Rate constant for cell development marker acquisition | (1/d) |
| $k_{nhh}$ | Rate constant for cell development marker acquisition | (1/d) |
| $x$ | Leakage factor | (-) |
| $a_m$ | For Langmuir function (monocytes) | (-) |
| $b_m$ | For Langmuir function (monocytes) | (-) |
| $a_H$ | For Langmuir function (HLA-DR) | (-) |
| $b_H$ | For Langmuir function (HLA-DR) | (-) |
| $b_{nhh}$ | For Langmuir function (mature neutrophils) | (-) |

## Results and discussion

Figure 3 shows the cluster centres after scaling and Table 3 gives average patient data for each cluster. Figure 4 gives mean day 90 survival versus the Simplified Acute Physiology (SAPSII) health assessment score, as well as the Trauma: Sepsis: Surgery fractions for each cluster. From Figure 4, the percentage D90 survival in each cluster is directly correlated to its sepsis fraction. Clusters 0 and 1 contain the patients with least severe illness. This is indicated by high survival rates and low values in their medical assessments. These being the SAPSII, Charlson comorbidity and Sequential Organ Failure Assessment (SOFA) scores. They also have the lowest rates of secondary infection and fewest sepsis patients. The difference between these two clusters is in neutrophil maturity. Cluster 1 is a group of 29 patients with relatively high CD16-, CD10+ neutrophils who resemble cluster 0 (123 patients) in all other respects. This suggests that the threshold for CD16-, CD10+ neutrophils used here does not quite match the boundary between a healthy or unhealthy response. The remaining three patient groups have severity of illness increasing in the order cluster 3 < 4 < 2, with cluster 2 patients having the lowest survival rates, highest SAPSII, Charlson and SOFA scores, the greatest chance of secondary infection and the highest proportion of sepsis patients.

**Figure 3.**
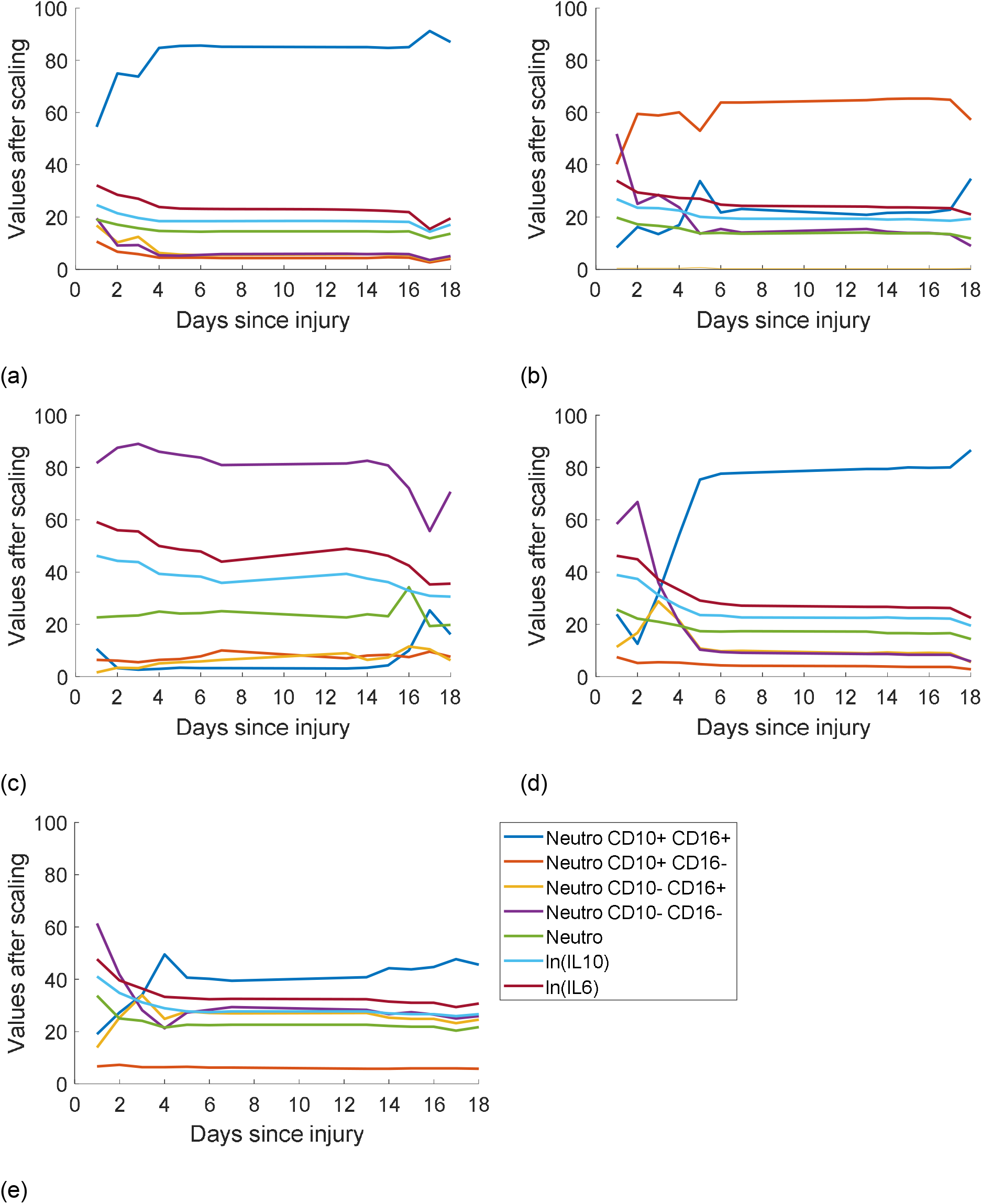
Cluster centres for neutrophil count, neutrophil phenotype fractions and logs of IL6 and IL10 after scaling for clusters 0 (a), 1 (b), 2 (c), 3 (d) and 4 (e)

**Figure 4.**
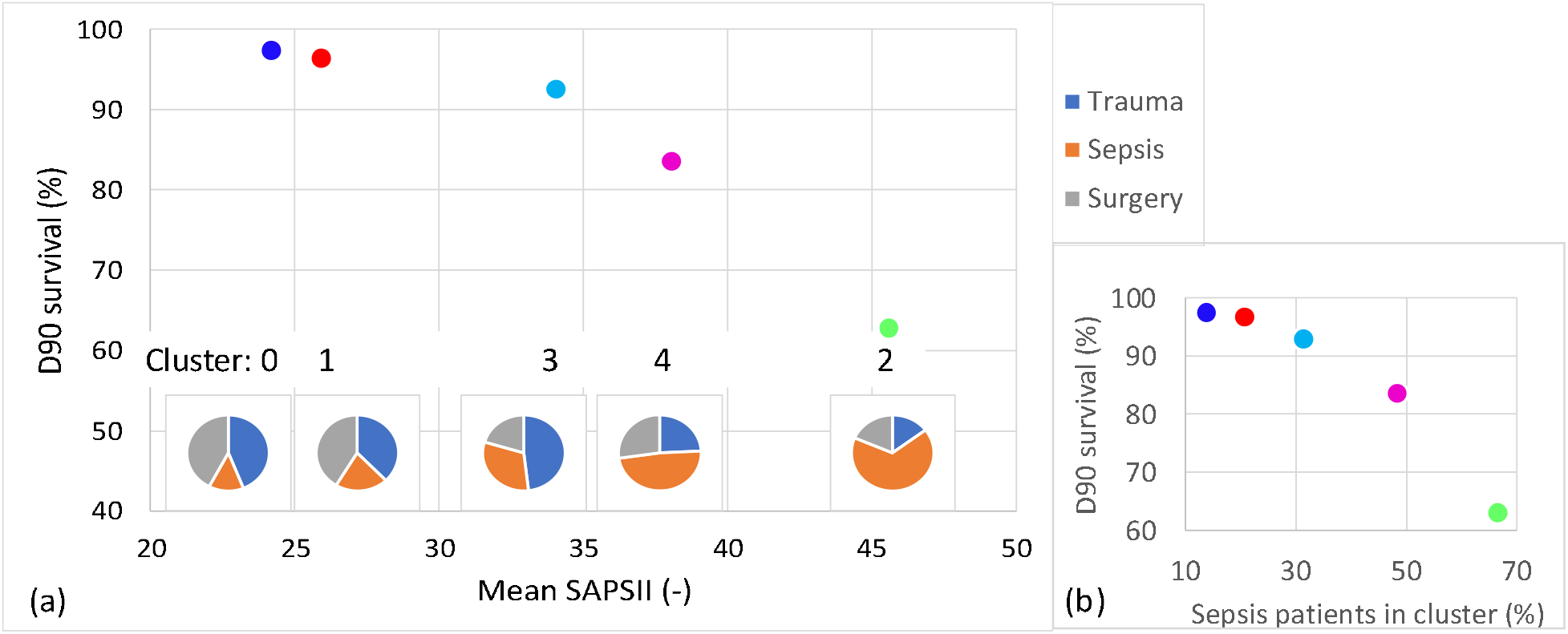
Distribution of clusters: mean day 90 survival vs SAPS II and Trauma: Sepsis: Surgery fractions for each cluster (a) and vs Sepsis patients in each cluster (b).

**Table 3.**
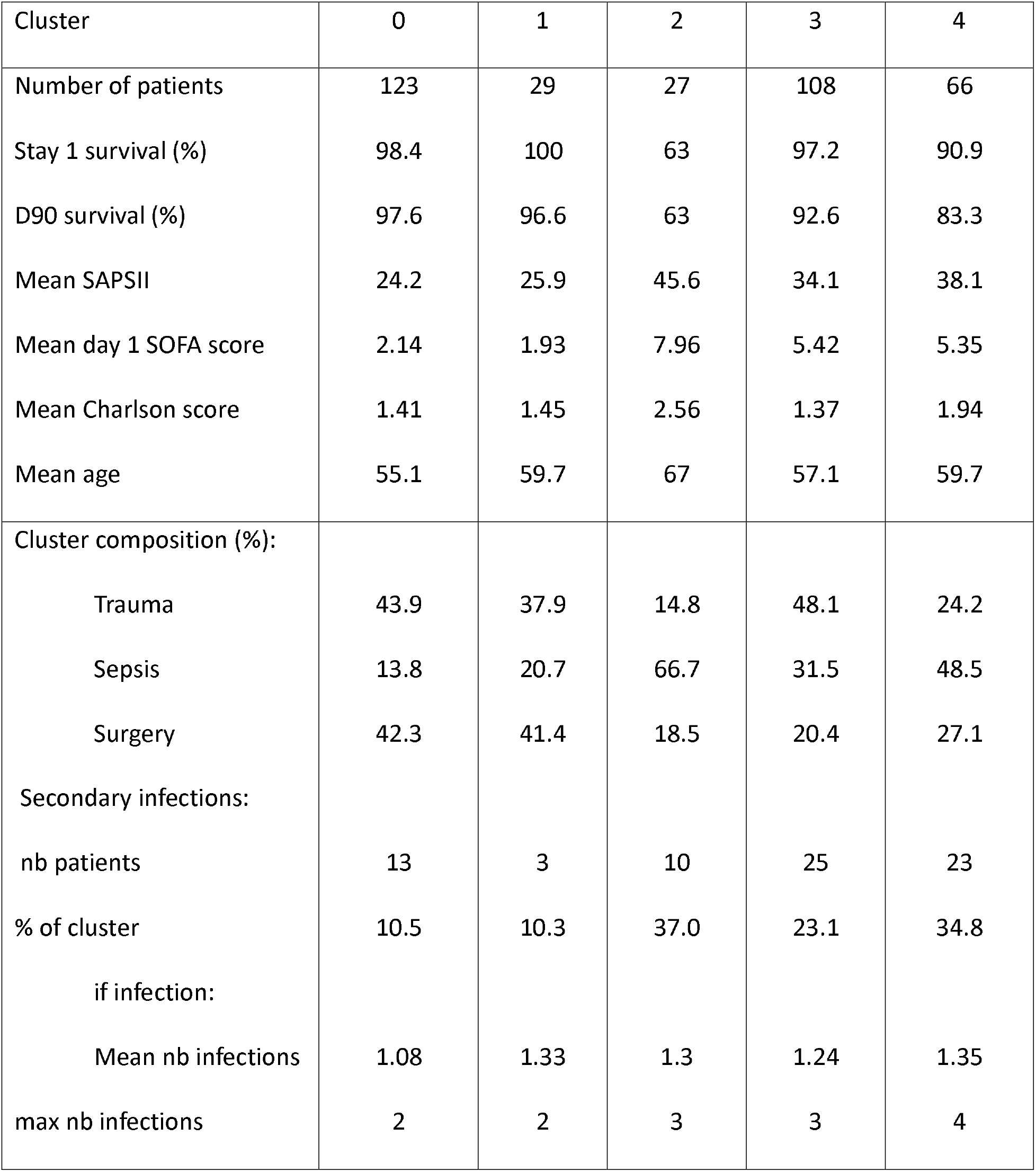
Average patient data for each cluster

| Cluster | 0 | 1 | 2 | 3 | 4 |
| --- | --- | --- | --- | --- | --- |
| Number of patients | 123 | 29 | 27 | 108 | 66 |
| Stay 1 survival (%) | 98.4 | 100 | 63 | 97.2 | 90.9 |
| D90 survival (%) | 97.6 | 96.6 | 63 | 92.6 | 83.3 |
| Mean SAPSII | 24.2 | 25.9 | 45.6 | 34.1 | 38.1 |
| Mean day 1 SOFA score | 2.14 | 1.93 | 7.96 | 5.42 | 5.35 |
| Mean Charlson score | 1.41 | 1.45 | 2.56 | 1.37 | 1.94 |
| Mean age | 55.1 | 59.7 | 67 | 57.1 | 59.7 |
| Cluster composition (%): |  |  |  |  |  |
| Trauma | 43.9 | 37.9 | 14.8 | 48.1 | 24.2 |
| Sepsis | 13.8 | 20.7 | 66.7 | 31.5 | 48.5 |
| Surgery | 42.3 | 41.4 | 18.5 | 20.4 | 27.1 |
| Secondary infections: |  |  |  |  |  |
| nb patients | 13 | 3 | 10 | 25 | 23 |
| % of cluster | 10.5 | 10.3 | 37.0 | 23.1 | 34.8 |
| if infection: |  |  |  |  |  |
| Mean nb infections | 1.08 | 1.33 | 1.3 | 1.24 | 1.35 |
| max nb infections | 2 | 2 | 3 | 3 | 4 |

The silhouette score^52,59^ for this clustering is 0.233 which indicates significant overlap between clusters. Examples of this can be seen in Figure 5, which compares IL6 data and CD16+, CD10+ neutrophil data for clusters 0 and 2, the two most different clusters. So, comparison of the different attributes for each group, including SAPSII score and number of secondary infections, shows an overlap, but the clusters are well spread across the spectrum of severity of illness and each cluster includes patients with every type of injury considered.

**Figure 5.**
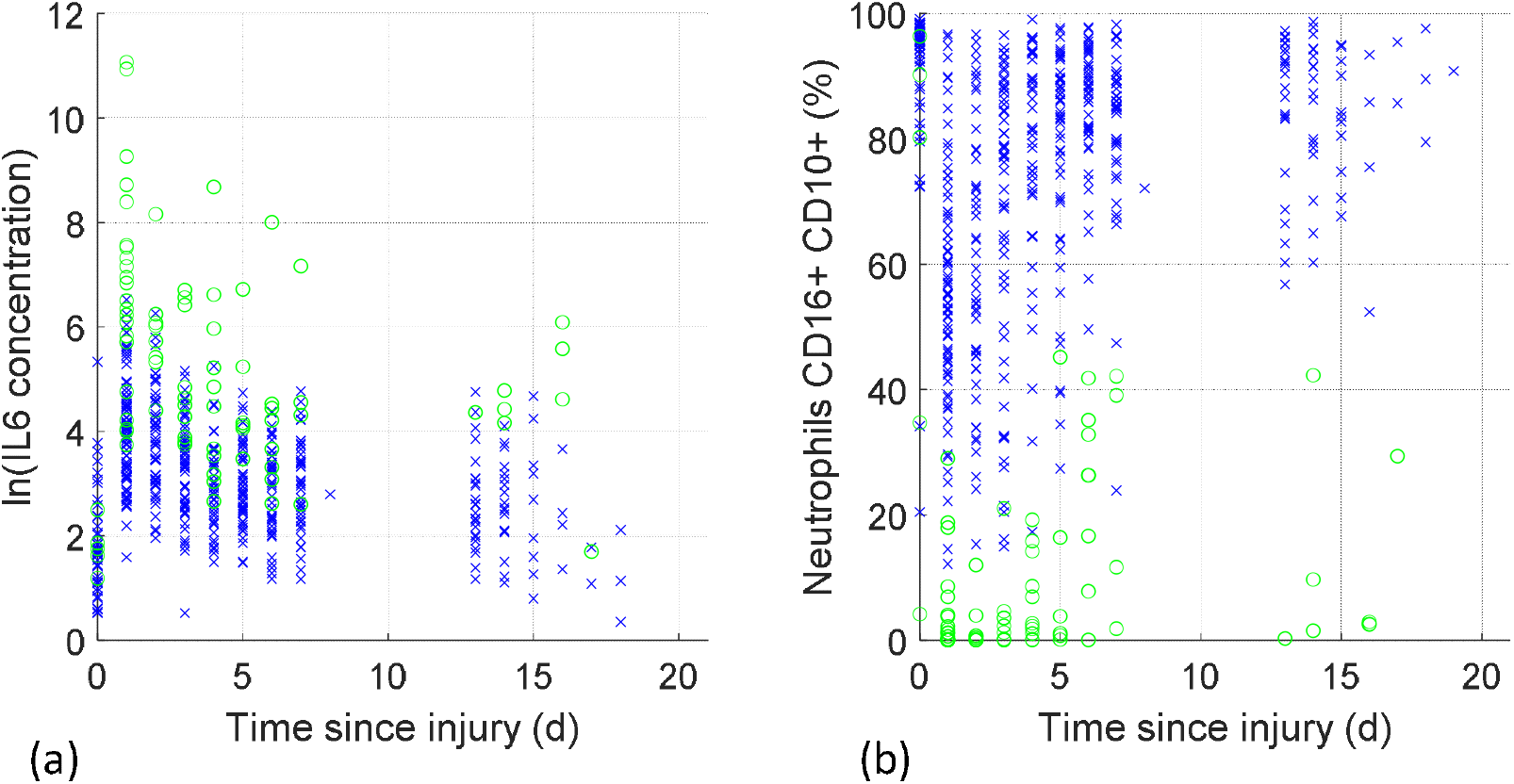
Clusters 0 (x) and 2 (o) datapoints for ln(IL6 concentration) (a) and CD16+, CD10+ neutrophils (b)

Figure 6 and Figure 7 show that the in-house model represents the IL6 and IL10 data well for all clusters. A few outlying very high data points were removed from clusters 1, 3 and 4 after day 30 where the patient response might have been due to something other than the initial injury. The cluster 0 IL10 results appear slightly over-fitted because the same model is always used, only the parameters vary. So, these IL6 and IL10 curves are used to calculate *F*. Clusters 0 and 1 have, by far, the lowest IL6 and IL10 levels and cluster 2 has the highest. Clusters 3 and 4 reach similar maximum IL6 concentrations. Whereas for IL10, the cluster 3 maximum is much higher than that of cluster 4 with a faster recovery of both IL6 and IL10. This is coherent as IL10 is anti-inflammatory and should therefore act to reduce IL6 concentration. Cluster 3 has the largest proportion of trauma victims, who are younger on average, and therefore likely to have a better immune response than cluster 4 patients.

**Figure 6.**
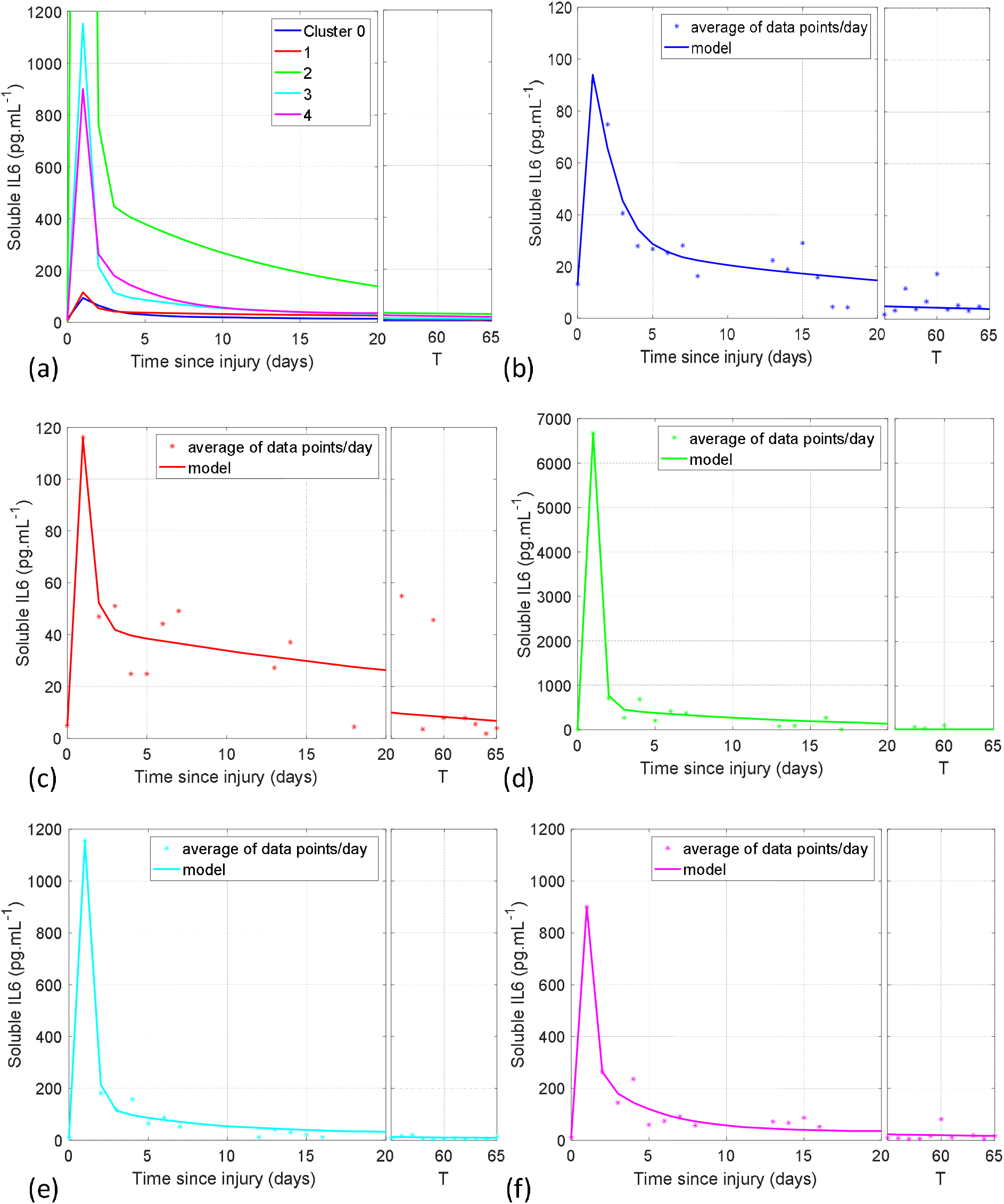
Calculated IL6 curves for all clusters (a) and comparison of calculated and measured daily average IL6 data for each cluster (b) to (f).

**Figure 7.**
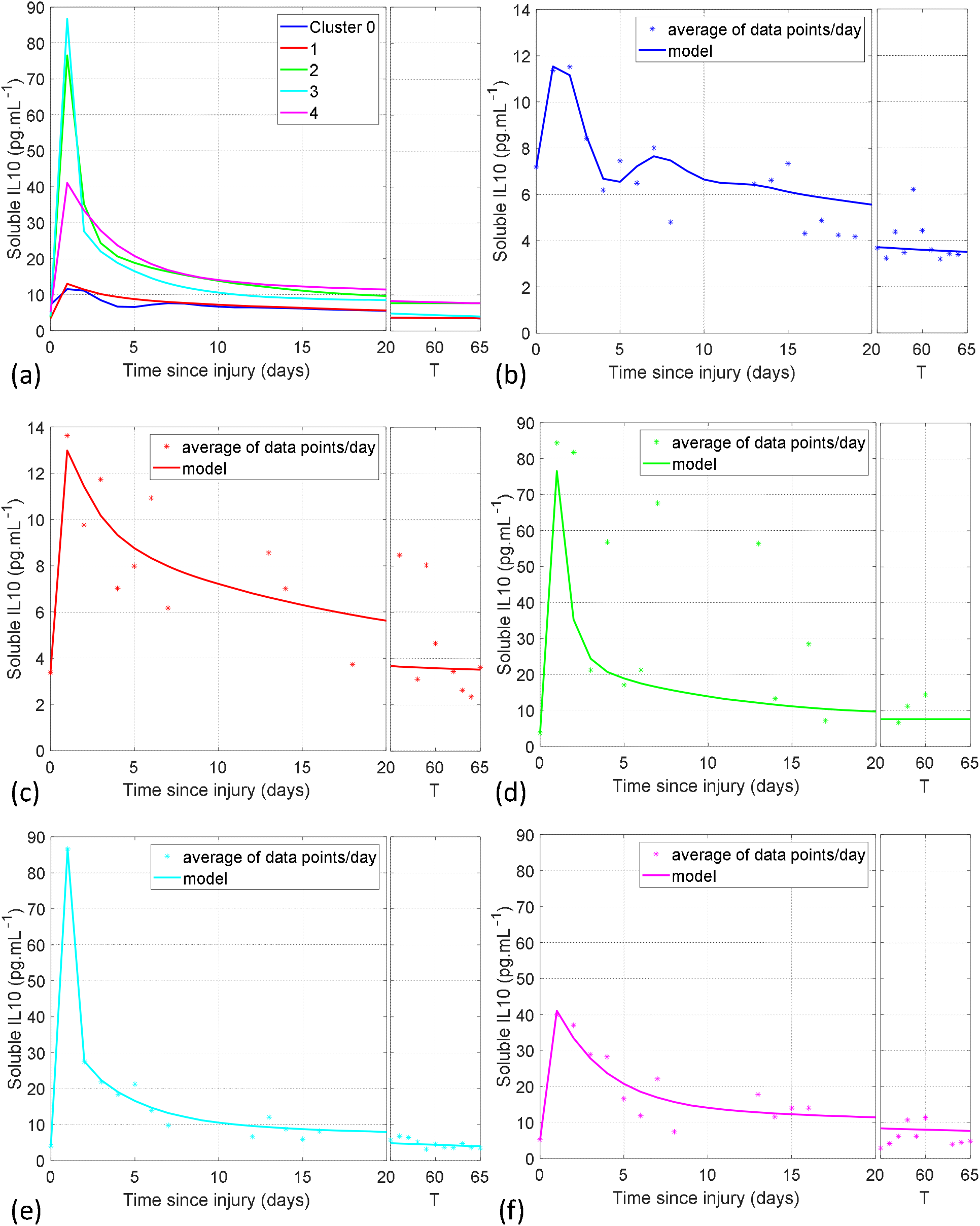
Calculated IL10 curves for all clusters (a) and comparison of calculated and measured daily average IL10 data for each cluster (b) to (f).

Figure 8 shows *F* for all the clusters. The shape of *F* closely resembles the IL6 curves, peaking on day 1, then declining rapidly at first, then more slowly. The ratio IL6/IL10 is always slightly greater than for the healthy controls meaning that *F* remains positive. *F* varies enormously in amplitude, with the day 1 maximum for cluster 2 almost 100 fold that of cluster 0. The initial, day zero, values of IL6 and IL10 are estimated from surgery patient data (there is no initial data for trauma or sepsis patients).

**Figure 8.**
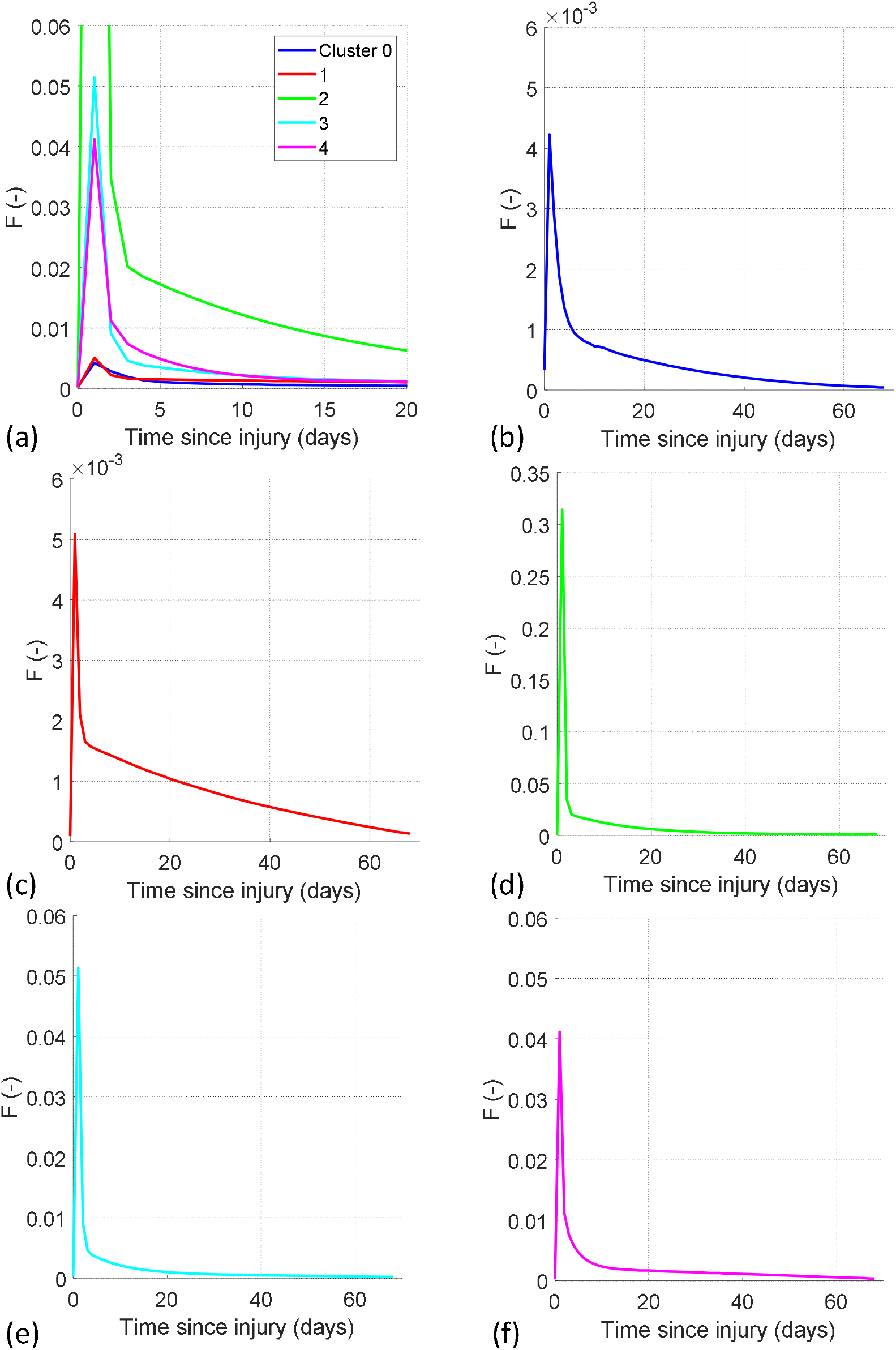
Calculated function, F, for all clusters (a) and clusters 0 (b), 1 (c), 2 (d), 3 (e) and 4 (f).

Table 4 gives rate constants *k_p,H_, k_nhl_, k_nhh_* and *k_nlh_* determined from the surgery patients’ pre-operation data. Discounting cluster 1, neutrophil maturation rate parameters calculated from the pre-operative surgery patients data, are lower for clusters 2 and 4, suggesting that the patients in most danger could already be identified at this early pre-surgery stage.

**Table 4.**
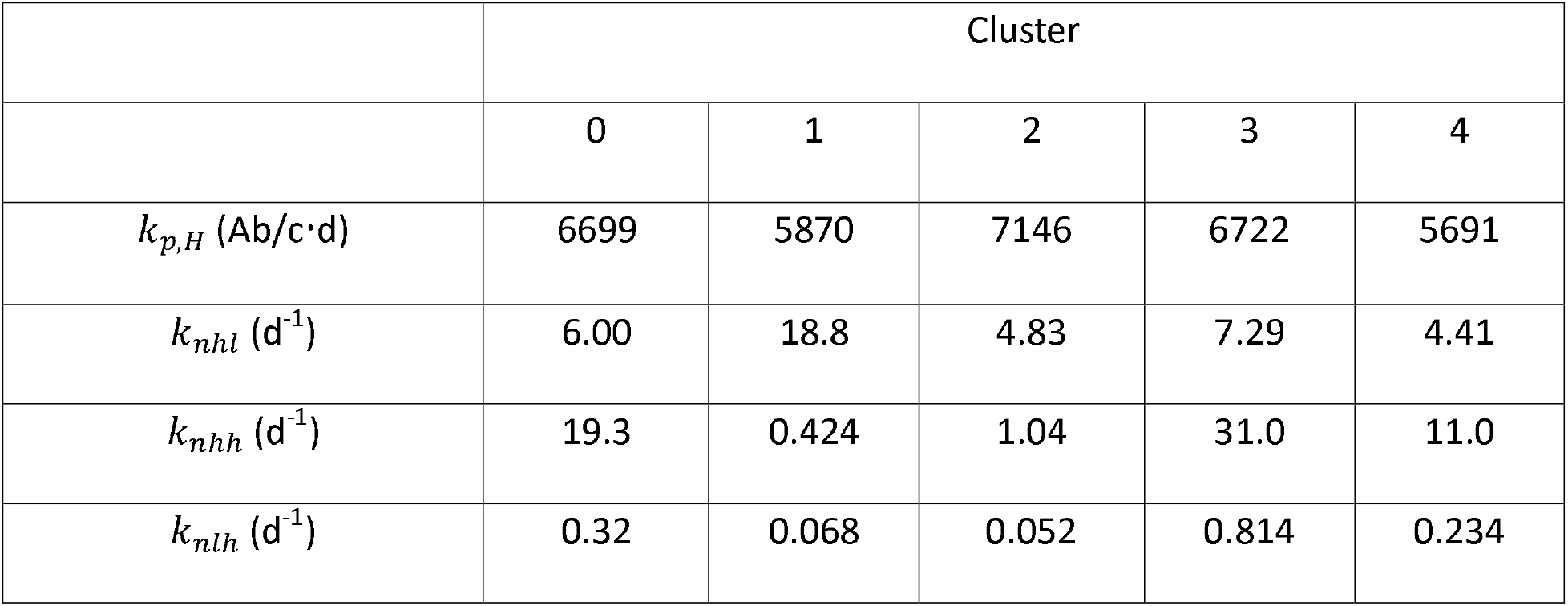
Parameter values calculated from surgery patients t0 data

|  | Cluster |  |  |  |  |
| --- | --- | --- | --- | --- | --- |
|  | 0 | 1 | 2 | 3 | 4 |
| $k_{p,H}$ (Ab/c·d) | 6699 | 5870 | 7146 | 6722 | 5691 |
| $k_{nhl}$ (d<sup>-1</sup>) | 6.00 | 18.8 | 4.83 | 7.29 | 4.41 |
| $k_{nhh}$ (d<sup>-1</sup>) | 19.3 | 0.424 | 1.04 | 31.0 | 11.0 |
| $k_{nlh}$ (d<sup>-1</sup>) | 0.32 | 0.068 | 0.052 | 0.814 | 0.234 |

Figure 9, Figure 10, Figure 11 and Figure 12 give model results for all clusters after parameter estimation. Figure 9 gives the measured and calculated average daily neutrophil counts plus and minus one standard deviation, with Figure 9 (a) the calculated neutrophil counts for each cluster over the first seven days after injury. Clusters 0 and 1 are almost identical, with a maximum on days 1 and 2 followed by a recovery. Clusters 3 and 4 have a similar dynamic with a higher maximum whilst cluster 2 peaks later and reduces much more slowly. There is quite a lot of variance in the data, confirming the overlapping clusters identified by the low silhouette score and demonstrated in Figure 5. This wide variance highlights the difficulty of identifying any individual sepsis, trauma or surgery patient trajectory from their initial condition and therefore the potential benefit of including dynamic data for a set of blood parameters. For visibility, similar data ranges are omitted from Figure 10, Figure 11 and Figure 12. The model fits the first week of data well for all the clusters. For the more severely ill patients in clusters 2, 3 and 4, there is quite a lot of scatter in the later data points and, since there are also fewer D60 blood samples spread over a relatively long time period, the confidence limits for the average values are quite wide. The scatter also makes sense because, by this time, some patients may be suffering from diverging consequences of the initial injury or unrelated health events. Cluster 2 stands out as having a slower dynamic over the whole time period. The model calculates a more rapid recovery of cluster 3 neutrophil count, compared to cluster 4, due to the relative IL6 and IL10 concentrations per Figure 6 (a) and Figure 7 (a).

**Figure 9.**
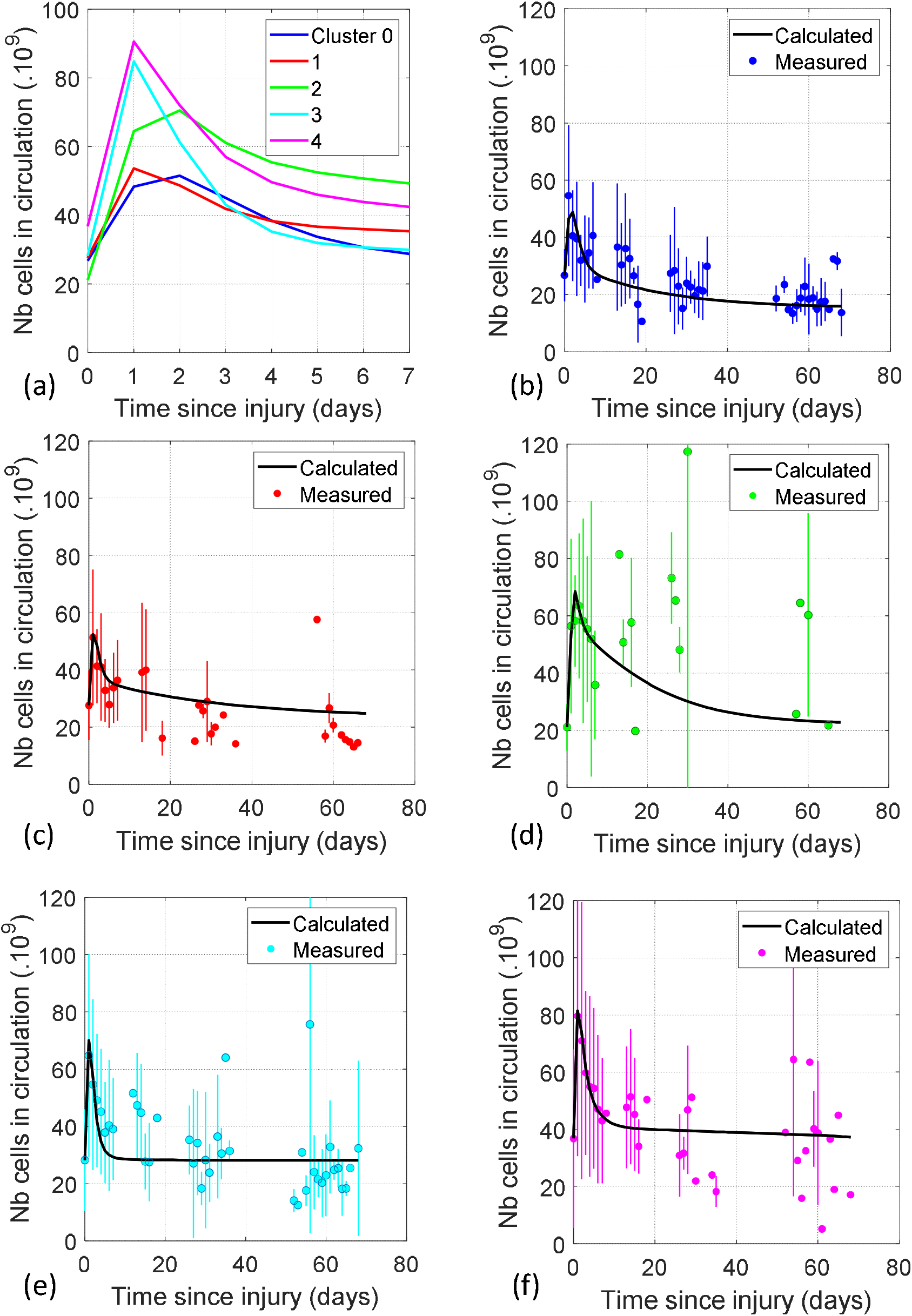
Calculated and measured average daily neutrophil counts ± one standard deviation for clusters all (a), 0 (b), 1 (c), 2 (d), 3 (e) and 4 (f).

**Figure 10.**
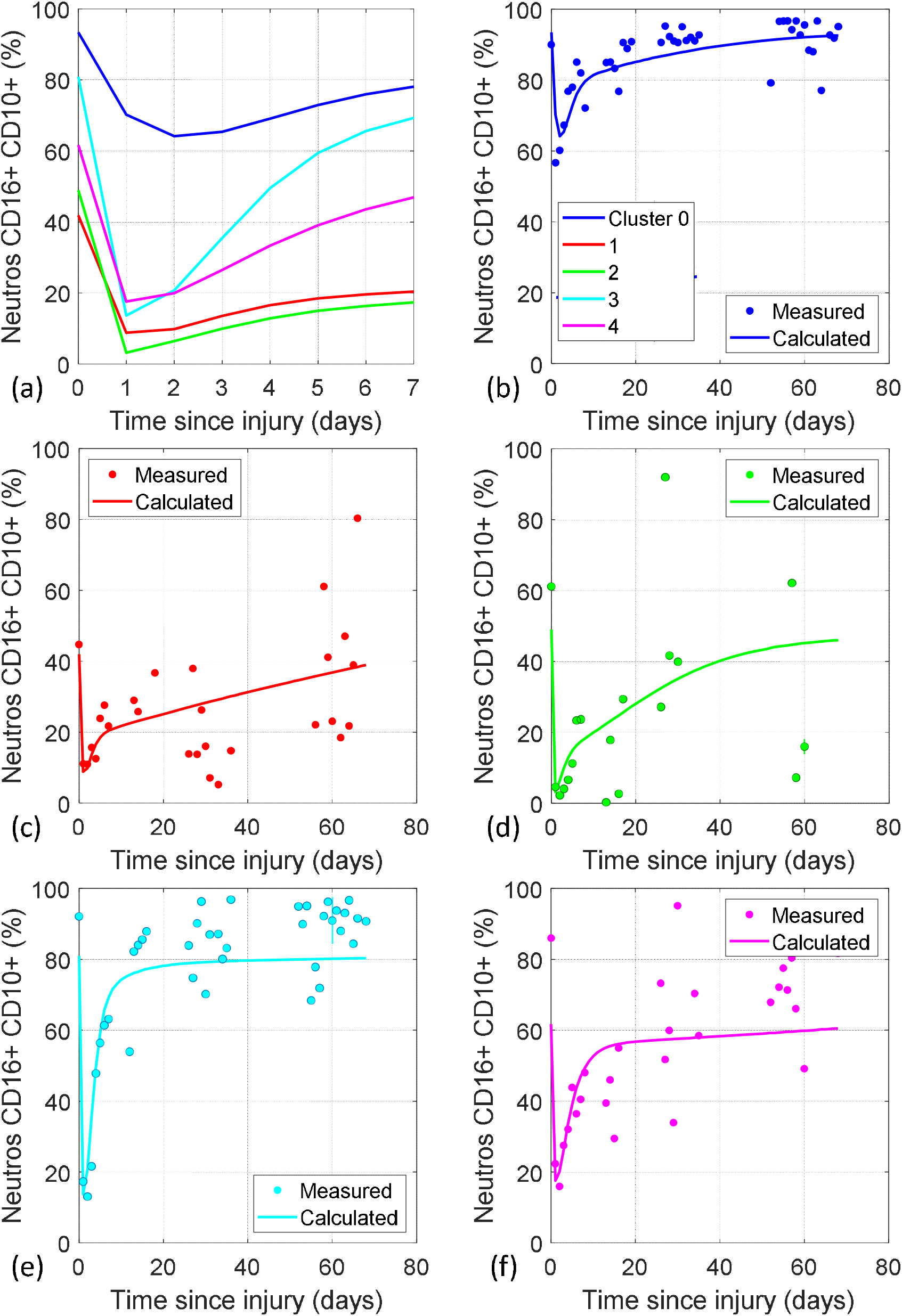
Calculated and measured average daily percent mature neutrophils for clusters all (a), 0 (b), 1 (c), 2 (d), 3 (e) and 4 (f).

**Figure 11.**
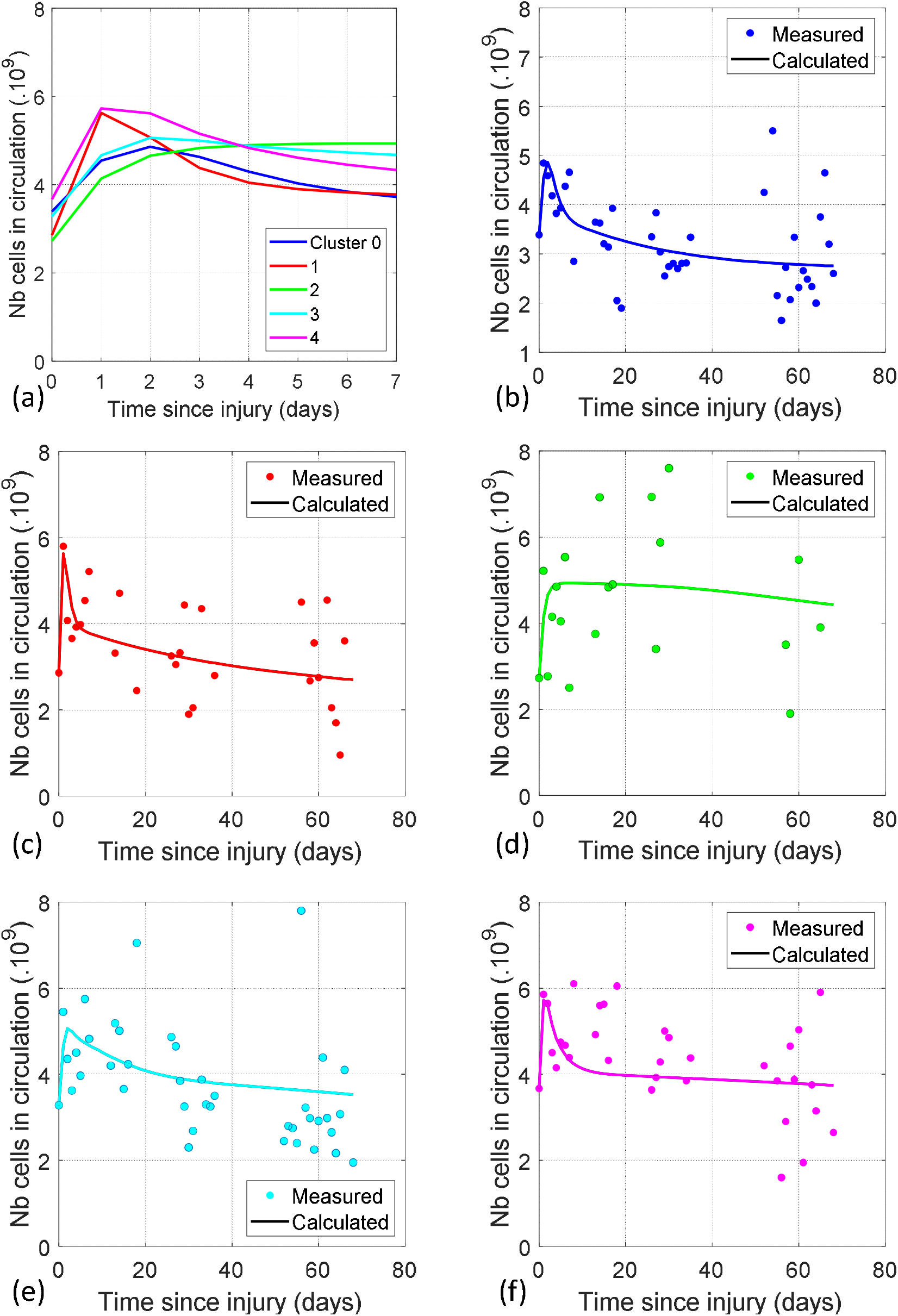
Calculated and measured average daily monocyte count for clusters all (a), 0 (b), 1 (c), 2 (d), 3 (e) and 4 (f).

**Figure 12.**
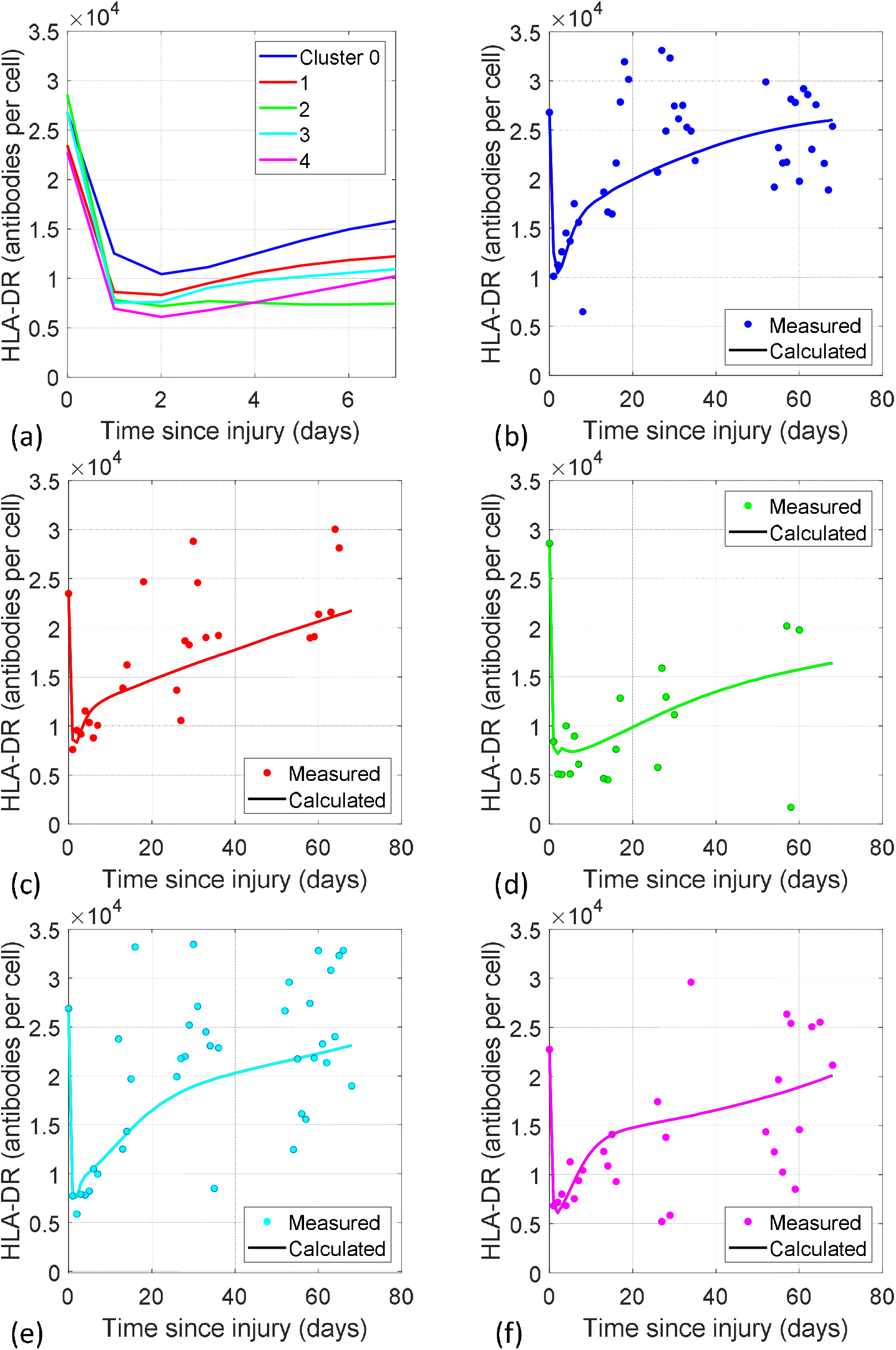
Calculated and measured average daily HLA-DR per monocyte for clusters all (a), 0 (b), 1 (c), 2 (d), 3 (e) and 4 (f).

Figure 10 gives the calculated percentage of CD16+ CD10+ neutrophils circulating and the average measured daily values for each cluster. Figure 10 (a) compares the results over the first seven days after injury. Except for cluster 1, the model predicts the order of severity of illness, 0 < 1< 3 < 4 < 2. The exception of cluster 1 is logical because this cluster resembles cluster 0, apart from the percentage of CD16+ CD10+ neutrophils (see Figure 1(a) and (b)). Again, cluster 2 has a different profile with the later data points quite scattered. The model picks up the trend in the CD16+ CD10+ neutrophils data. However, it doesn’t quite capture the full variation. This could be because of the assumed distribution of the neutrophil bone marrow and blood counts, which are not well understood, particularly the quantity and dynamics of marginated neutrophils^26,60^.

Figure 11 gives the results for the monocyte count. There is wide variance in the data but the model fits the early average data points quite well for all the clusters. Again, the slower dynamic of cluster 2 is visible in the calculated results for the first seven days and over the whole time period.

Figure 12 gives the calculated and measured mHLA-DR for each cluster. The model fits the initial data points very well. By day 4 the calculated HLA-DR levels for each cluster are in the same order as the severity of illness: 0 < 1< 3 < 4 < 2. However, whilst clusters 0, 1, 3 and 4 follow a similar dynamic, with a minimum on day 1 or 2, cluster 2 does not recover at all during the first week. The mHLA-DR quantity is associated with the level of immunosuppression, with 10 – 15K Ab/c described as moderate and <5 or <8K Ab/c severe^14^. By this measure, all clusters are immunosuppressed throughout the first week after injury. Initially, clusters 2, 3 and 4 are severely immunosuppressed and the cluster 2 recovery does not achieve a healthy level. The patterns for mHLA-DR expression kinetics in sepsis by Leijte et al.^46^ and Bodinier et al.^47^ were identified as high expressors, early improvers, delayed improvers and decliners. Here, the corresponding clusters would be 0/1, 3, 4 and 2 respectively.

In the following we analyse the parameter estimation results and the calculated impact of the inflammation function. There are eleven estimated parameters as listed previously in Table 2. Two Eq. (3) parameters each for neutrophils, monocytes and HLA-DR, one for mature neutrophils, three reaction rate constants and the leakage parameter. Estimated rate constants for each cluster are presented to three significant figures in Table 5 with their confidence levels. Correlation coefficients are given in Figure 13 and plots of 90 day survival rates against estimated parameter values are shown in Figure 14. The cluster 1 parameters have low confidence levels. This is the small group of 29 patients whose evolution is like cluster 0 except for the percentage of CD16-CD10+ neutrophils. There is sometimes correlation between parameters, *a* and *b*, particularly for the less severe clusters.

**Table 5.**
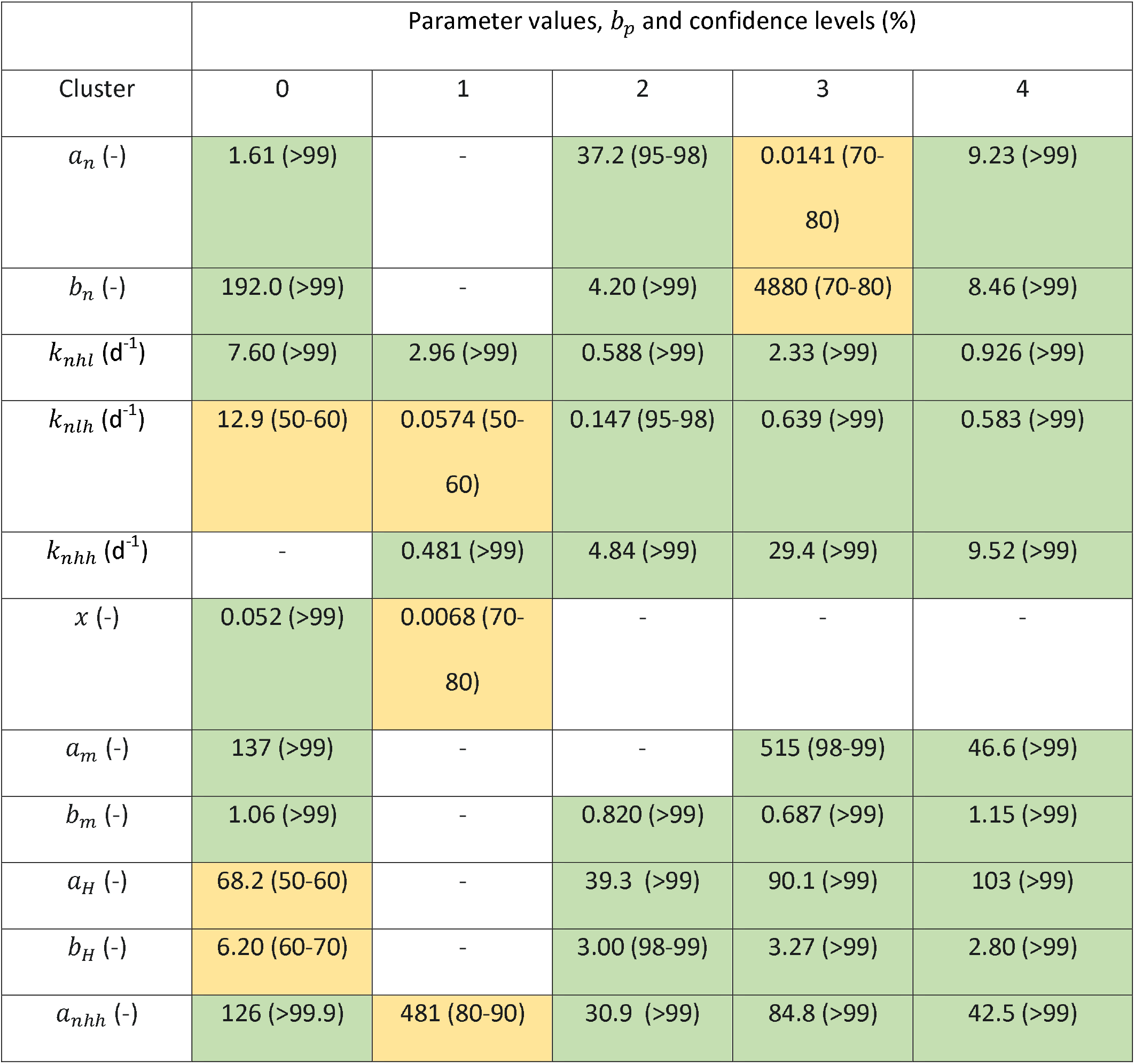
Estimated parameter values and their confidence levels for each cluster: yellow 50-90%, green >90 confidence.

**Figure 13.**
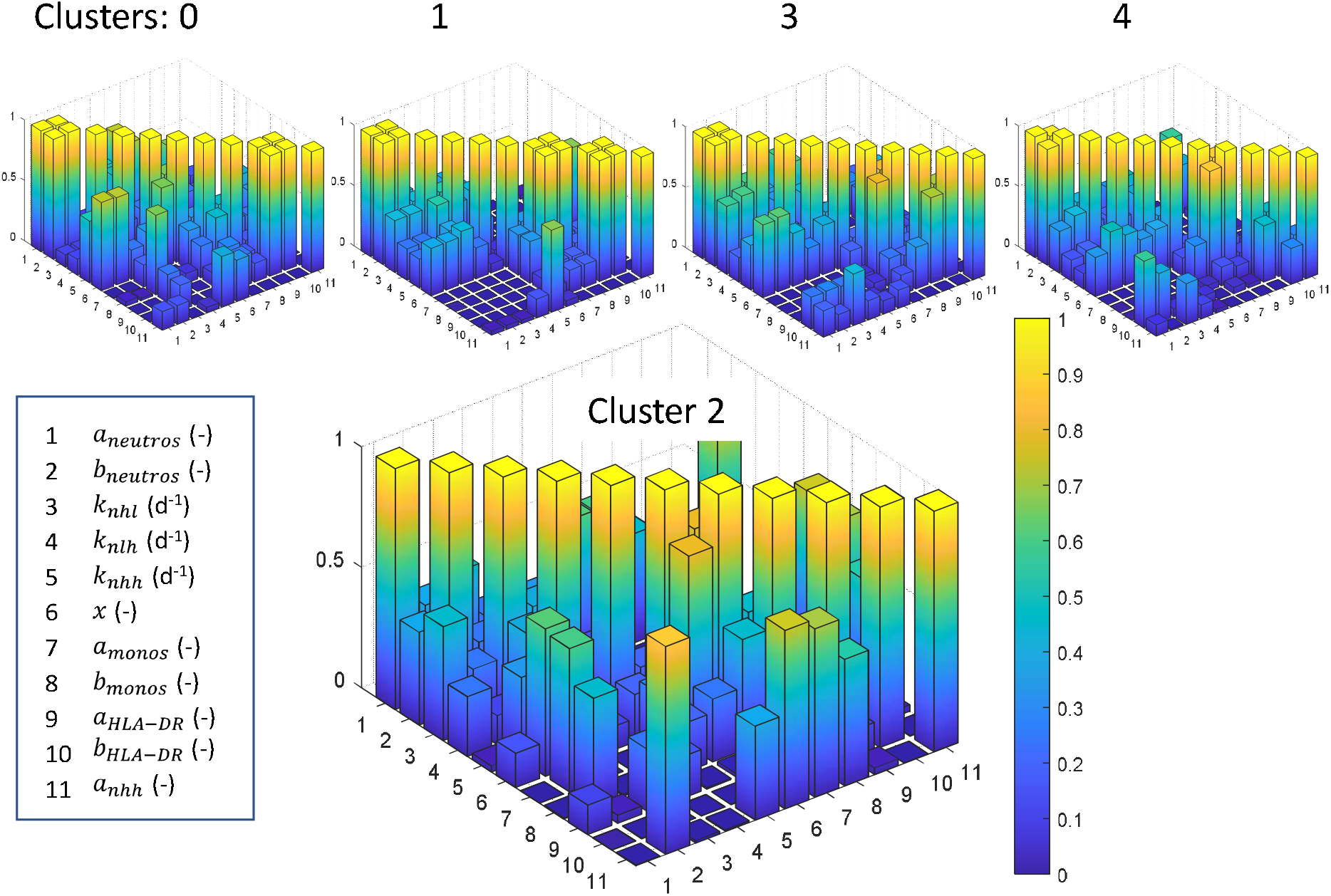
Correlation between estimated parameters for each cluster.

**Figure 14.**
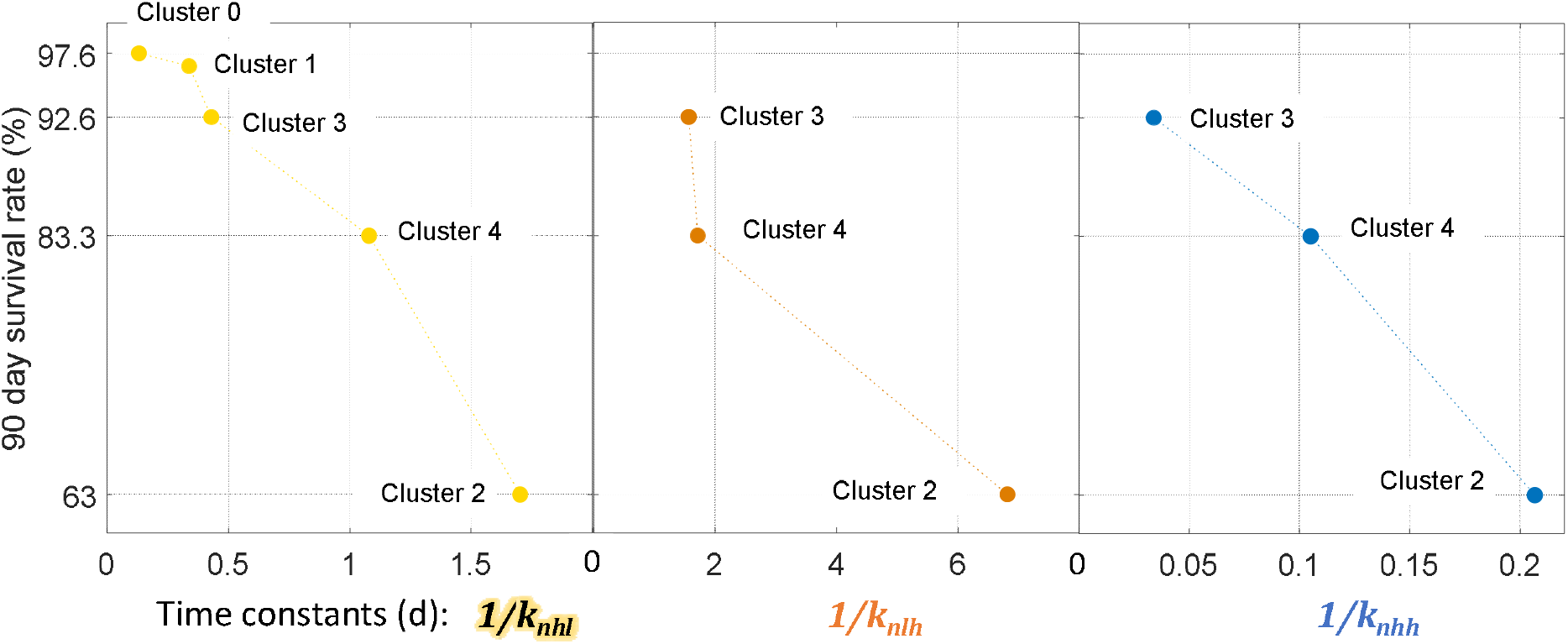
Plots of statistically significant estimated rate constants for all clusters..

Most of the estimated neutrophil maturation rate constants, *k_n_*, are statistically significant and they tend to decrease with severity of illness, with those for cluster 2 particularly low. The initial parameter values were calculated from the pre-surgery patients and the estimation has adjusted *k_nlh_* and *k_nhh_* slightly. However, the estimated maturation rate constants for addition of CD16, *k_nhl_* are generally much lower than the calculated pre-surgery values, suggesting that this cell maturation process could be reduced after injury. Figure 14 shows the neutrophil maturation time constants for each cluster and maturation step. The estimated parameters also show that addition of CD10 is more rapid for CD16+ neutrophils, as might be expected to occur in healthy individuals and suggesting that, if CD16 expression is retarded, so is CD10 addition. The maximum residence time in compartments 2 and 3, with no inflammation, is two days. So, by this model, the whole cluster 2 neutrophil population could not achieve maturity, even with no inflammation. In line with this, it has been suggested elsewhere that CD10-neutrophil levels could be an indicator for patients at risk from of excessive inflammatory immune response^61^.

The leakage parameter is significant for clusters 0 and 1 but not for the more severely affected clusters. The parameter *a_nhh_* varies with severity of illness, so that the more severely ill, the greater the ratio of *F* to neutrophil disappearance rate.

Considering the Eq. (3) parameters, *a* and *b*, for neutrophils, monocytes and mHLA-DR. Eq. (3) translates *F* into its response, *f*. As an example, *F* and *f*, for clusters 3 and 2 are shown in Figure 15. When the curves for *F* and *f* are similar, as for cluster 3 neutrophil count (Figure 15 (a)), one parameter is sufficient to describe the linear relation, *a* and *b* are correlated and have very low statistical significance. This is also the case for the CD16+ CD10+ neutrophil response in all cases. On the other hand, for cluster 2 neutrophils illustration, the shapes of the curves for *F* and *f* are quite different (Figure 15 (b)), suggesting a limiting factor. This could be a lack of available cytokine receptors at the cell surface or affinity between receptors and cytokines, for example.

**Figure 15.**
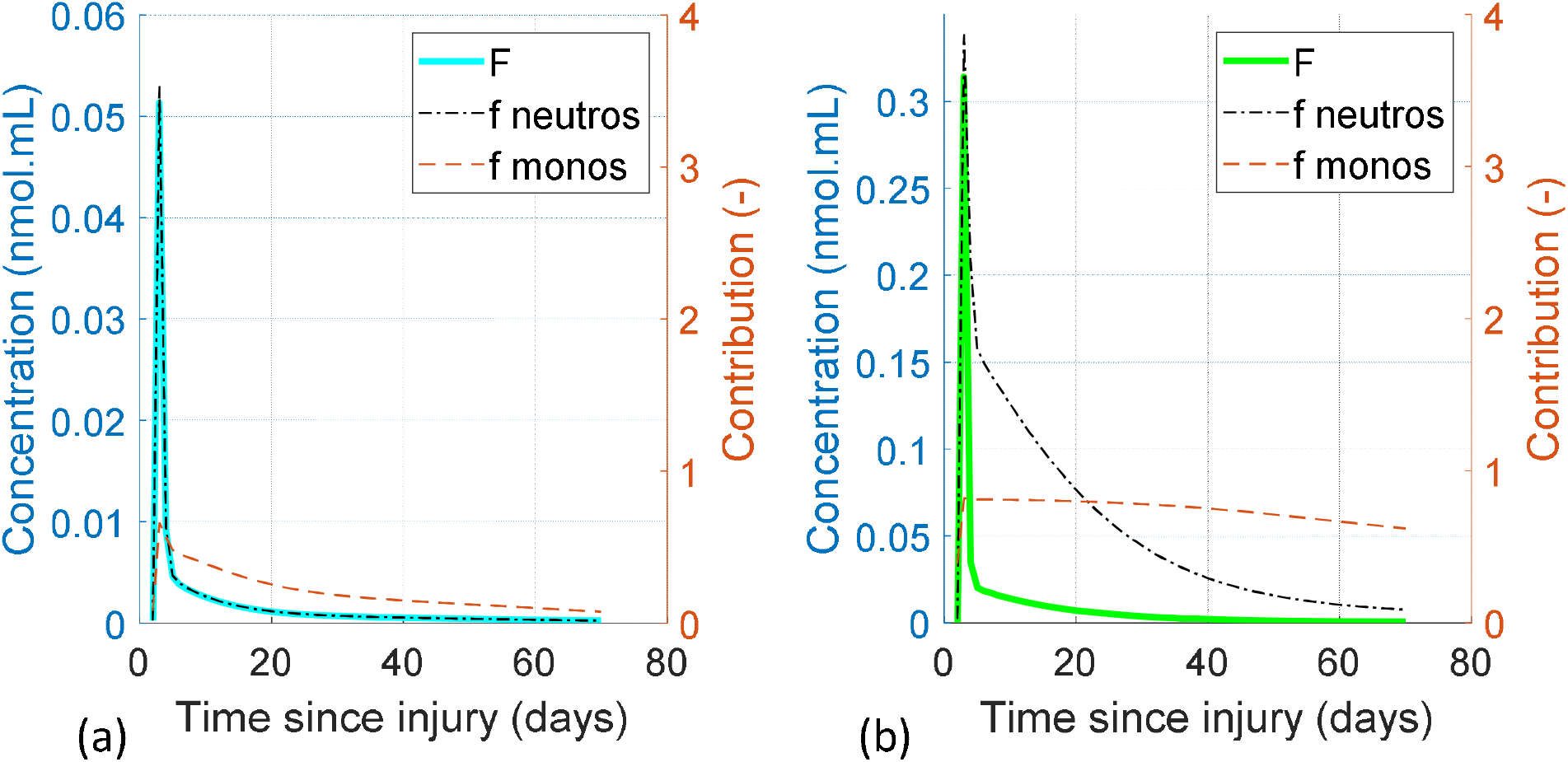
Response to inflammation function for neutrophils and monocytes for cluster 3 (a) and cluster 2 (b).

## Conclusion

To sum up, the time-series clustering has allowed sepsis, trauma and surgery patients to be grouped into clusters across the spectrum of severity of illness. Clusters 0 and 1 are the least severe, in terms of all the parameters considered here, and are very similar except for the CD16+ CD10+ neutrophil level, suggesting that four clusters might provide a better description and care should be taken in using neutrophil maturity as a tool for diagnosis. Cluster 2 contains the most severely ill group of patients, with the greatest perturbation of each measure and the highest risk. Clusters 3 and 4 are intermediate and reach a similar peak level of IL6 but cluster 3 has relatively more IL10 on day 1, faster recovery and better outcomes. Clusters 3, 4 and 2 are comparable with the early improvers, delayed improvers and decliners suggested by Leijte et al.^46^.

The dynamic model gives a good representation of the neutrophil count. The model doesn’t quite capture the extremes of the CD16+ CD10+ neutrophil profiles, possibly due to the assumptions on their distribution between the bone marrow and blood populations. Representation of the mHLA-DR profiles is good.

In healthy individuals, *F* is very small and the pre-operative values for the surgery patients and the final values of *F* are slightly positive for each cluster. This demonstrates that homeostasis is not tied to a single value of *F* but, more likely, a distribution of low values. Under the conditions immediately after injury, IL6 and IL10 levels are high and *F* works well. *F* is always positive, suggesting that the immunosuppression after sepsis and trauma derives, not from a negative signal, but from the cells’ capacity to respond. The function *f* takes into account the state of the cells (quantity of surface receptors and affinity for their associated cytokines) and the capacity of the host to respond to *F* and by producing healthy cells.

Dynamic modelling has also shown that all the patient clusters can be represented by the same model using different parameters, although some aspects of the model are redundant for some groups of patients. Despite the data set extending to 68 days after injury, the model is based only on the expected behaviour of the innate immune system. The commonality of the immediate response to injury^62^ makes this straightforward. A more detailed model would capture the impact of the adaptive immune response on later data points. However, the data becomes more scattered with time as more parameters come into play, such as the adaptive immune system or other health events and the data becomes more disparate. It is interesting that the pre-operative neutrophil data for the surgery patients identifies the clusters at greatest risk, suggesting that these patients might be identified before undergoing surgery. The parameter estimation results suggest that the neutrophils and monocytes of cluster 2, the most seriously ill, are at the limit of their responsiveness.

Potential future work includes study of the variability within each cluster via the range of individual responses.

## Acknowledgements

The authors gratefully thank all members of the REALISM study group:

– HCL: Sophie ARNAL, Caroline AUGRIS-MATHIEU, Frédérique BAYLE, Liana CARUSO, Charles-Eric BER, Asma BEN-AMOR, Anne-Sophie BELLOCQ, Farida BENATIR, Anne BERTIN-MAGHIT, Marc BERTIN-MAGHIT, André BOIBIEUX, Yves BOUFFARD, Jean-Christophe CEJKA, Valérie CERRO, Jullien CROZON-CLAUZEL, Julien DAVIDSON, Sophie DEBORD-PEGUET, Benjamin DELWARDE, Robert DELEAT-BESSON, Claire DELSUC, Bertrand DEVIGNE, Laure FAYOLLE-PIVOT, Alexandre FAURE, Bernard FLOCCARD, Julie GATEL, Charline GENIN, Thibaut GIRARDOT, Arnaud GREGOIRE, Baptiste HENGY, Laetitia HURIAUX, Catherine JADAUD, Alain LEPAPE, Véronique LERAY, Anne-Claire LUKASZEWICZ, Guillaume MARCOTTE, Olivier MARTIN, Marie MATRAY, Delphine MAUCORT-BOULCH, Pascal MEURET, Céline MONARD, Florent MORICEAU, Guillaume MONNERET, Nathalie PANEL, Najia RAHALI, Thomas RIMMELE, Cyrille TRUC, Thomas UBERTI, Hélène VALLIN, Fabienne VENET, Sylvie TISSOT, Abbès ZADAM
– bioMérieux: Sophie BLEIN, Karen BRENGEL-PESCE, Elisabeth CERRATO, Valérie CHEYNET, Emmanuelle GALLET-GORIUS, Audrey GUICHARD, Camille JOURDAN, Natacha KOENIG, François MALLET, Boris MEUNIER, Virginie MOUCADEL, Marine MOMMERT, Guy ORIOL, Alexandre PACHOT, Estelle PERONNET, Claire SCHREVEL, Olivier TABONE, Julien TEXTORIS, Javier YUGUEROS MARCOS
– BIOASTER: Jérémie BECKER, Frédéric BEQUET, Yacine BOUNAB, Florian BRAJON, Bertrand CANARD, Muriel COLLUS, Nathalie GARCON, Irène GORSE, Cyril GUYARD, Fabien LAVOCAT, Philippe LEISSNER, Karen LOUIS, Maxime MISTRETTA, Jeanne MORINIERE, Yoann MOUSCAZ, Laura NOAILLES, Magali PERRET, Frédéric REYNIER, Cindy RIFFAUD, Mary-Luz ROL, Nicolas SAPAY, Trang TRAN, Christophe VEDRINE
– SANOFI: Christophe CARRE, Pierre CORTEZ, Aymeric DE MONFORT, Karine FLORIN, Laurent FRAISSE, Isabelle FUGIER, Sandrine PAYRARD, Annick PELERAUX, Laurence QUEMENEUR
– ESPCI: Andrew GRIFFITHS, Stephanie TOETSCH
– GSK: Teri ASHTON, Peter J. GOUGH, Scott B. BERGER, David GARDINER, Iain GILLESPIE, Aidan MACNAMARA, Aparna RAYCHAUDHURI, Rob SMYLIE, Lionel TAN, Craig TIPPLE

## Notes

### Competing Interest Statement

The authors have declared no competing interest.

